# Molecular investigations on a chimeric strain of *Staphylococcus aureus* sequence type 80

**DOI:** 10.1101/2020.04.08.031591

**Authors:** Darius Gawlik, Antje Ruppelt-Lorz, Elke Müller, Annett Reißig, Helmut Hotzel, Sascha D. Braun, Bo Söderquist, Albrecht Ziegler-Cordts, Ralf Ehricht, Stefan Monecke

## Abstract

An Eritrean patient was admitted with suspected tuberculous cervical lymphadenitis. While no mycobacteria were detected in pus from this process, culture yielded PVL-positive, methicillin-susceptible *Staphylococcus aureus*. Microarray hybridisation assigned the isolate to clonal complex (CC) 80 but revealed unusual features, including the presence of the ORF-CM14 enterotoxin homologue and of an ACME-III element as well as the absence of *etD* and *edinB.* The isolate was subjected to both, Illumina and Nanopore sequencing allowing characterisation of deviating regions within the strain’s genome. Atypical features of this strain were attributable to the presence of two genomic regions that originated from other *S. aureus* lineages and that comprised, respectively, 3% and 1.4% of the genome. One deviating region extended from *walJ* to *sirB*. It comprised ORF-CM14 and the ACME-III element. A homologous, but larger fragment was also found in an atypical *S. aureus* CC1/ST567 strain whose lineage might have served as donor of this genomic region. This region itself is a chimera comprising fragments from CC1 as well as fragments of unknown origin. The other region of another 3% of the genome comprised the region from *htsB* to *ecfA2*. It was very similar to CC1 sequences. This suggests either an incorporation of CC1 DNA into the study strain, or it might alternatively suggest a recombination event affecting “canonical” CC80. As the study strain bears witness of several recombination events, such complex and large-scale events cannot be rare and exceptional, despite a mainly clonal nature of *S. aureus*. Although the exact mechanism is not yet clear, chimerism seems to be an additional pathway in the evolution of *S. aureus*, possibly being responsible for the transmission also of virulence and resistance factors. An organism that can shuffle, swap or exchange major parts of its genome by a yet unknown mechanism would have an evolutionary advantage compared to a strictly clonal organism.

## Introduction

*Staphylococcus aureus* (*S. aureus*) is a versatile pathogen that colonises or infects a large fraction of the world’s human population as well as several species of wild and domestic animals. Thus, it can asymptomatically colonise its carriers, or alternatively cause various infections ranging from superficial skin and soft tissue infections to serious bacteremia including infective endocarditis. Many of its virulence factors are variably present and their genes are localized on mobile genetic elements such as plasmids, phages, transposons or on pathogenicity islands. In recent decades, some strains of *S. aureus* acquired resistance to many or most antibiotics. Again, resistance genes are localized on mobile, or potentially mobile, genetic elements such as staphylococcal chromosomal cassette *mec* (SCC*mec*) cassettes. Despite a vast variety of variable, mobile elements, and despite some incremental, mutation-driven variation, the overall structure of the *S. aureus* genome is conservative with all core genomic elements being present in all strains in one uniform sequential arrangement. Multilocus sequence typing (MLST) enables the unambiguous assignment of isolates to taxonomic units, sequence types (ST) and clonal complexes (CC), based on numbered alleles of seven housekeeping genes assuming that these genes cannot be lost or truncated because of their crucial function and that the accumulation of mutations in their sequences is purely a function of time. This lead to a model of a clonal evolution of the *S. aureus* core genome that is driven by a time-dependent accumulation of single point mutations allowing classification based on a few marker genes into a limited number of clonal complexes comprising a number of sequence types that differ only in random mutations in these marker genes as well as of others.

However, some *S. aureus* strains show features that cannot be explained neither by accumulation of single point mutations nor by acquisition or loss of mobile genetic elements. For instance, there is a Russian CC8 strain in which nearly half of the genome is inverted [1]. Other strains show evidence of large-scale recombination events, with considerably large fragments of their genomes originating from other lineages and being inserted at the appropriate position of the recipient strain. Such a phenomenon was first postulated for ST239 strains, in which deviant alleles of *arcC* and *spa, aur, clfB* and *isaB*, of the capsule type (5 instead of 8) and the presence of *cna* indicate an integration of a CC30 DNA fragment of approximately 635,000 base pairs (or ca. 20% of the genome) into a CC8 recipient strain, with the integration site being localised around *oriC* [2]. Another strain, ST2249, harbours ST239 DNA comprising fragments of both, CC8 and CC30 that is integrated into a CC45 genome [3]. Further examples for chimeric strains are ST34 and ST42 (in which CC10 fragments are integrated into CC30 genomes) [2] or CC398 strains that harbour fragments of CC9 origin [4, 5]. Such observations indicate that large scale recombination events played a role in driving the evolution of *S. aureus* but the underlying molecular mechanisms are not yet described. In the present paper, we describe another, new chimeric strain that comprises a backbone of CC80 genomic DNA and two separate large inserts that attracted attention because of a presence of ORF-CM14 and an absence of *edinB* and *etD*.

*S. aureus* CC80 is a lineage that is commonly associated with recurrent and/or severe skin and soft tissue infections (SSTI) since a majority of isolates carries the phage-borne Panton-Valentine leukocidin (PVL), a virulence factor associated with SSTI. One PVL-positive, methicillin-resistant CC80 strain, with a SCC*mec* IVc element, is sporadically found in Western Europe [6-14] while it is widespread in Mediterranean countries including Greece [15, 16], Turkey [17], Lebanon [18], Malta [19], Tunisia [20, 21] and Algeria [22-24] as well as in the Middle East/Arabian Gulf regions [25-28]. It can also commonly be found in European travellers to these regions [10, 17]. Methicillin-susceptible CC80 strains are uncommon but geographically widespread in Africa [29-31] from where this lineage originated [14].

The isolate described herein was initially subjected to microarray hybridisation, primarily for typing and detection resistance and toxin genes. The procedure revealed unusual features that could be explained by a large-scale horizontal gene transfer. This observation prompted further investigations including Illumina and Nanopore sequencing in order to map the entire genome of the isolate and a search for the donor strain of regions assumed to be introduced by the gene transfer.

## Material and Methods

### Clinical background and isolates

An Eritrean patient was admitted in 2015 to the Dresden University Hospital (Dresden, Saxony, Germany) with a cervical skin and soft tissue infection that originally was suspected to be suppurative tuberculous lymphadenitis. While no mycobacteria were detected (neither immediately by microscopy after staining for acid-fast bacilli nor subsequently in MGIT and Ogawa cultures), culture of pus from this process yielded a PVL-positive, methicillin-susceptible *S. aureus* (ANR570100).

A second isolate was further characterised because of certain similarities with the study isolate (see below). It was isolated in 2002. It originated from an approximately 50 years old female patient with lobar pneumonia probably secondary to an influenza B infection. She was a Swedish citizen and denied any traveling outside Sweden.

### Microarray-based typing

The *S. aureus* isolates were initially characterized using different DNA microarray-based assays. Probes, primers as well as amplification and hybridization protocols have previously been described in detail [32-34].

### Illumina sequencing

Genomic DNA for whole-genome sequencing was prepared from culture on Columbia blood agar incubated overnight at 37°C. DNA was prepared using the Qiagen DNA isolation kit (Qiagen, Hilden, Germany) according to manufacturer’s instructions after an enzymatic lysis step with lysostaphin, lysozyme and RNAse as previously described [32-34]. Afterwards, whole-genome sequencing was carried out using the Illumina HiSeq2500 genome analyser (Illumina HiSeq 2500 platform, Illumina, Essex, UK). The reads were assembled to contigs using SPAdes.

Sequencing of the two strains was performed at two geographically distant facilities and at different dates (Jena, Germany, and Örebro, Sweden, in spring and autumn 2018, respectively), ruling out any possibility of carry-over contaminations.

### Nanopore sequencing

The Nanopore Oxford MinION platform was used for whole-genome sequencing. Briefly, no size selection was performed and the DNA library was generated using the native barcoding expansion kit EXP-NBD103 and the Nanopore sequencing kit SQK-LSK109 (Oxford Nanopore Technologies, Oxford, UK) according to manufacturer’s instructions. The used flow cell FLO-MIN106 (R9-Version) was primed by the flow cell priming kit EXP-FLP001 (Oxford Nanopore). The protocol “1D Native barcoding genomic DNA” was used in version NBE_9065_v109_revB_23May2018 (Last update: 03/09/2018). The albacore basecaller (Oxford Nanopore) translated the minion raw data (FAST5) into short quality tagged sequence reads (FASTQ). After barcode trimming using Poreshop (https://github.com/rrwick/Porechop, release v0.2.4), canu (https://github.com/marbl/canu, release v1.7.1) was used to assemble the short reads. After nano-polishing (https://github.com/jts/nanopolish, release v0.11.3), the corrected sequence data were used for a direct comparison to the Illumina sequence data (see below).

### Bioinformatic analysis

Iterated BLAST searches were used for analysis of individual contigs in this genome (https://blast.ncbi.nlm.nih.gov/Blast.cgi). This analysis was conducted using automated scripts for full text comparison and BLAST analysis and an in-house database of known, annotated and previously identified *S. aureus* genomes, genes and fragments to the query sequence. This enables the determination of identity, gene content, clonal parentage and of position within the genome of each contig given the constant order of core genomic genes in *S. aureus*.

Contigs were compared to the CC80 strain 11819-97, GenBank CP003194.1/SAMN02603886. This is a PVL-positive strain with a SCC*mec* IV element and – as essentially all canonical CC80 strains – with an *etD/edinB* pathogenicity island. It can be regarded as representative for CC80. Its genome has a size of 2,846,546 nt and an average G/C content of 32.9%. They were also compared to the long-known CC8 strain COL, GenBank CP000046, and to MW2, BA000033, as reference sequence for CC1.

Finally, Nanopore and Illumina sequences were aligned manually for reasons discussed below.

## Results

### Comparison of methods

A total of 36 Illumina contigs was considered to be chromosomal. Another one was assumed to contain plasmid-borne sequences (including *blaZ* and *cadX*; see below). The average fragment site of the library was 220 nt. Visual inspection and comparison of these contigs to the Nanopore sequences revealed faulty assemblies of four contigs that needed to be split into two “sub-contigs” each in order to allow alignment to the Nanopore sequence. Most significantly, Illumina/velvet failed to resolve a ca. 5,000 nt region within the ACME-III element that consisted of repetitive sequences. On the other hand, Nanopore showed a poor resolution of poly-A and poly-T sequence fragments resulting in the loss of approximately 15,800 nucleotides across the entire chromosome.

### Characterisation of the clinical isolate

Array hybridization revealed the presence of the enterotoxin homologue ORF-CM14 and of an ACME-III element, as well as the absence of *edinB* and *etD*. Otherwise, the isolate matched previously characterized CC80 strains (see Supplemental file 1). In order to explain these discrepancies, it was sequenced using both, Illumina and Nanopore methods and resulting sequences were aligned resulting in a continuous chromosome with a total length of 2,789,663 nt and an overall G/C content of 32.98%. MLST was performed based on the consensus genome sequence and it yielded ST-80 (*arcC*-1, *aroE*-3, *glpF*-1, *gmk*-14, *pta*-11, *tpi*-51, *yqiL*-10).

A comparison of core chromosomal genes revealed that two regions in ANR570100 differed from CP003194. When visually inspecting the mapping of the ANR570100 reads to CP003194, in those two regions we were able to identify the true extend of these deviant regions (Figure 1).

**Figure 1:**
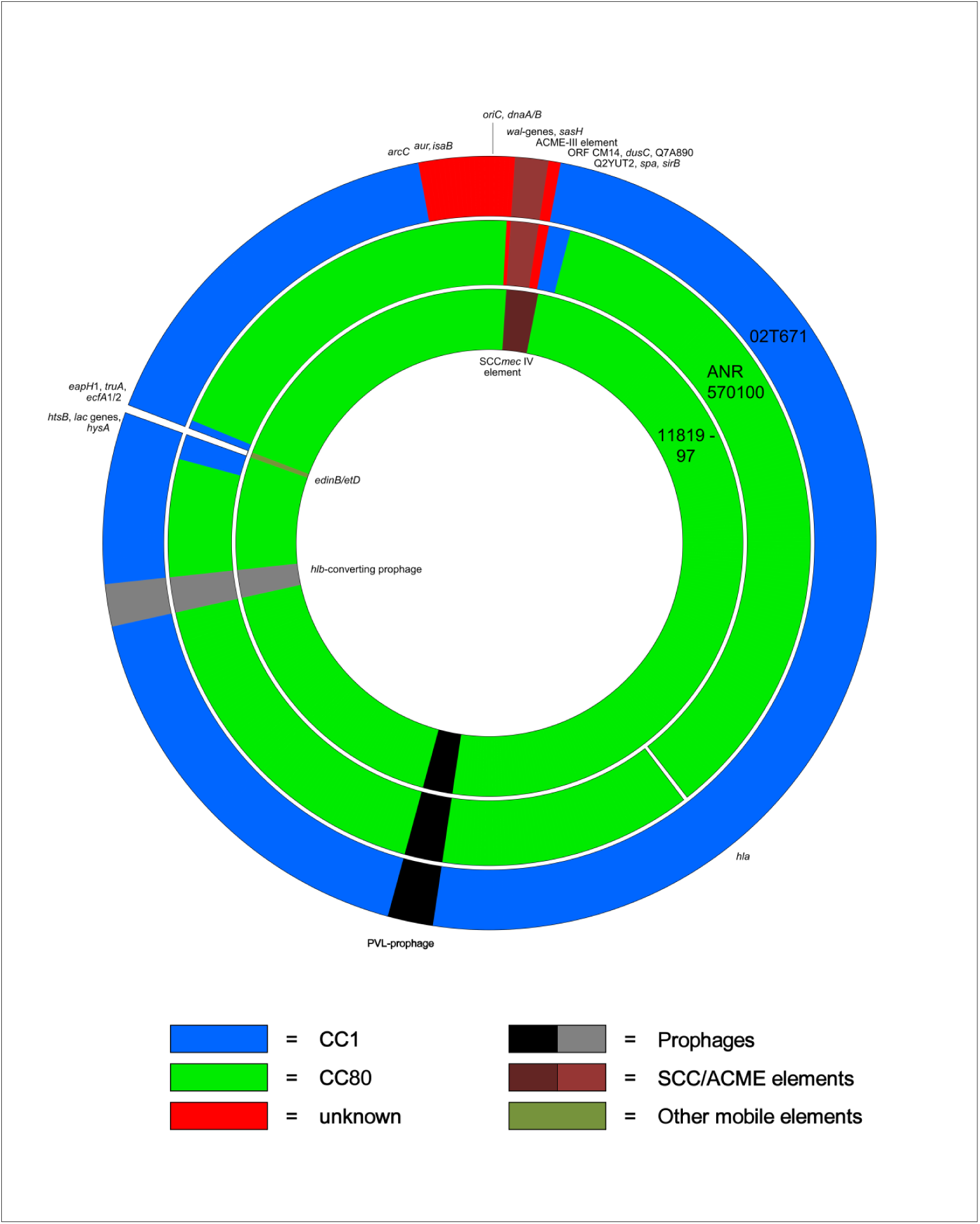
Schematic diagram of the genomes of 02T567 (outer circle), ANR570100 (middle circle) and the reference genome CC80, 11819-97, GenBank CP003194.1 (inner circle). Genomic fragments are colour-coded depending on their origin.

### Deviating Region 1

Deviating Region 1 extends from *walJ* (locus tag MS7_0024 in CC80, CP003194, or respectively SACOL0023 in CC8, CP000046) with a putative recombination sites located in the intergenic region between *walL* and *walJ*. It extends to certainly include *sirB* (MS7_0106, SACOL0098), possibly even to *sbnE* (MS7_0112, SACOL0104) although the differences to canonical CC80 sequences are not large enough to clearly determine a recombination site. It can be estimated at 84,363 nt (based on a consensus of the Illumina and the Nanopore sequences, and including *walJ* to *sirB*). This corresponds to roughly 3% of the genome and includes *ca.* 34,000 nt belonging to the ACME-element.

The gene content of Deviating Region 1 is described in Table 1 where it is also compared to a CC80 reference sequence CP003194 as well as to 02T-671.

**Table 1:** Genes in Deviating Region 1 in comparison to canonical CC80, to 02T-671 and to canonical CC1.

|  | Comparison of ANR570100 to CC80 reference sequence 11819-97 (CP003194) | Comparison of ANR570100 to 02T-671 | Comparison of 02T-671 to MW2 | Provenance in ANR570100 | Provenance in 02T-671 |
| --- | --- | --- | --- | --- | --- |
| *walR* | 0 mismatches/702 bases (0%) | 2 mismatches/702 bases (0.28%) | 4 mismatches/702 bases (0.57%) | CC80 | unknown |
| *walK* | 5 mismatches/1827 bases (0.27%) | 12 mismatches/1827 bases (0.66%) | 14 mismatches/1827 bases (0.77%) | CC80 | unknown |
| *walH* | 18 mismatches/1335 bases (1.35%) | 36 mismatches/1335 bases (2.7%) | 36 mismatches/1335 bases (2.7%) | CC80 | unknown |
| *walI* | 0 mismatches/789 bases (0%) | 19 mismatches/789 bases (2.41%) | 19 mismatches/789 bases (2.41%) | CC80 | unknown |
| *walJ* | 17 mismatches/801 bases (2.12%) | 0 mismatches/801 bases (0%) | 18 mismatches/801 bases (2.25%) | unknown | unknown |
| *sasH* | 95 mismatches/2319 bases (4.1%) | 0 mismatches/2319 bases (0%) | 65 mismatches/2319 bases (2.8%) | unknown | unknown |
| *orfX* | 16 mismatches/480 bases (3.33%) | 0 mismatches/480 bases (0%) | 14 mismatches/480 bases (2.92%) | unknown | unknown |
| **ACME-III (see Table 2)** | *Not present in 11819-97 (which has SCCmec IVc instead)* | Identical element (see text and Table 3) | *Not present in MW2 (which has SCCmec IVa instead)* | ACME-III | ACME-III |
| **Q6GKL1** | *Not present in 11819-97/in canonical CC80* | 0 mismatches/576 bases (0%) | *Not present in MW2/in canonical CC1* | unknown | unknown |
| **Q6GKL6** | *Not present in 11819-97/in canonical CC80* | 0 mismatches/159 bases (0%) | *Not present in MW2/in canonical CC1* | unknown | unknown |
| **ORF-CM14** | *Not present in 11819-97/in canonical CC80* | 0 mismatches/780 bases (0%) | *Not present in MW2/in canonical CC1* | unknown | unknown |
| *dusC* | 29 mismatches/987 bases (2.94%) | 0 mismatches/987 bases (0%) | 26 mismatches/987 bases (2.63%) | unknown | unknown |
| **A6TXM6** | 23 mismatches/204 bases (11.27%) | 0 mismatches/204 bases (0%) | 25 mismatches/204 bases (12.25%) | unknown | unknown |
| **A6QD71** | 31 mismatches/297 bases (10.44%) | 0 mismatches/297 bases (0%) | 32 mismatches/297 bases (10.77%) | unknown | unknown |
| **Q5HJT2** | *Absent from ANR570100 and canonical CC80* | *Absent from ANR570100 and 02T-671* | *Absent from 02T-671 but present in canonical CC1* | unknown | unknown |
| **Q6GD34** | *Absent from ANR570100 and canonical CC80* | *Absent from ANR570100 and 02T-671* | *Absent from 02T-671 but present in canonical CC1* | unknown | unknown |
| **A6QD75** | *Absent from ANR570100 and canonical CC80* | *Absent from ANR570100 and 02T-671* | *Absent from 02T-671 but present in canonical CC1* | unknown | unknown |
| **A6QD76** | *Absent from ANR570100 and canonical CC80* | *Absent from ANR570100 and 02T-671* | *Absent from 02T-671 but present in canonical CC1* | unknown | unknown |
| **A8YZ18** | *Absent from ANR570100 and canonical CC80* | *Absent from ANR570100 and 02T-671* | *Absent from 02T-671 but present in canonical CC1* | unknown | unknown |
| **Q6GKK6** | 22 mismatches/612 bases (3.6%) | 1 mismatches/612 bases (0.16%) | 17 mismatches/612 bases (2.78%) | unknown | unknown |
| **Q7A890** | *Not present in 11819-97* | 1 mismatches/3153 bases (0.03%) | 104 mismatches/3153 bases (3.3%) | unknown | unknown |
| **Q2YUT2** | 1 mismatches/483 bases (0.21%) | 0 mismatches/483 bases (0%) | 2 mismatches/483 bases (0.41%) | CC1 or CC80 | CC1 |
| *plc* | 26 mismatches/987 bases (2.63%) | 0 mismatches/987 bases (0%) | 1 mismatches/987 bases (0.1%) | CC1 | CC1 |
| ***lpl*-SAOUHSC_00052** | 17 mismatches/771 bases (2.2%) | 0 mismatches/771 bases (0%) | 1 mismatches/771 bases (0.13%) | CC1 | CC1 |
| ***lpl*-SAOUHSC_00053** | 97 mismatches/771 bases (12.58%) | 0 mismatches/771 bases (0%) | 1 mismatches/771 bases (0.13%) | CC1 | CC1 |
| ***lpl*-MW0073** | 100 mismatches/693 bases (14.43%) | 0 mismatches/693 bases (0%) | 0 mismatches/693 bases (0%) | CC1 | CC1 |
| ***lipC3*-MW0074** | 42 mismatches/1377 bases (3.05%) | 0 mismatches/1377 bases (0%) | 0 mismatches/1377 bases (0%) | CC1 | CC1 |
| **Q8NYT6** | 17 mismatches/2238 bases (0.76%) | 1 mismatches/2238 bases (0.04%) | 20 mismatches/2238 bases (0.89%) | CC1 | CC1 ? |
| **Teg15as** | 0 mismatches/244 bases (0%) | 0 mismatches/244 bases (0%) | 0 mismatches/244 bases (0%) | *Related in all lineages in question* | *Related in all lineages in question* |
| **Q8NYT5** | 23 mismatches/1179 bases (1.95%) | 0 mismatches/1179 bases (0%) | 1 mismatches/1179 bases (0.08%) | CC1 | CC1 |
| ***norC*** | 27 mismatches/1389 bases (1.94%) | 0 mismatches/1389 bases (0%) | 0 mismatches/1389 bases (0%) | CC1 | CC1 |
| ***nptA*** | 14 mismatches/1662 bases (0.84%) | 0 mismatches/1662 bases (0%) | 0 mismatches/1389 bases (0%) | CC1 | CC1 |
| **Q2YUS5** | 17 mismatches/1776 bases (0.96%) | 0 mismatches/1776 bases (0%) | 1 mismatches/1776 bases (0.06%) | CC1 | CC1 |
| **DUF1648** | 0 mismatches/474 bases (0%) | 0 mismatches/474 bases (0%) | 0 mismatches/474 bases (0%) | *Related in all lineages in question* | *Related in all lineages in question* |
| ***lctP*-locus1** | 20 mismatches/1593 bases (1.26%) | 0 mismatches/1593 bases (0%) | 0 mismatches/1593 bases (0%) | CC1 | CC1 |
| ***txbi_lctP*** | 1 mismatches/72 bases (1.39%) | 0 mismatches/72 bases (0%) | 0 mismatches/72 bases (0%) | CC1 | CC1 |
| ***txbi_proteinA*** | 1 mismatches/62 bases (1.61%) | 0 mismatches/62 bases (0%) | 0 mismatches/62 bases (0%) | CC1 | CC1 |
| ***spa*** | ANR570100: *spa* t 1849<br>**07-23-34-33-13**<br>11819-97: *spa* t044<br>**07-23-12-34-34-33-34** | ANR570100: *spa* t 1849<br>**07-23-34-33-13**<br>02T671: *spa* t1242<br>**07-23-12-34-34-16-34-33-13** | 02T671: *spa* t1242<br>**07-23-12-34-34-16-34-33-13**<br>MW2: *spa* t128<br>**07-23-23-21-16-34-33-13** | *Related in all lineages in question* | *Related in all lineages in question* |
| ***tx_sarS*** | 0 mismatches/63 bases (0%) | 0 mismatches/63 bases (0%) | 0 mismatches/63 bases (0%) | *Related in all lineages in question* | *Related in all lineages in question* |
| ***sarS*** | 5 mismatches/753 bases (0.66%) | 0 mismatches/753 bases (0%) | 0 mismatches/753 bases (0%) | CC1 | CC1 |
| ***sirC*** | 7 mismatches/999 bases (0.7%) | 0 mismatches/999 bases (0%) | 0 mismatches/999 bases (0%) | CC1 | CC1 |
| ***sirB*** | 4 mismatches/996 bases (0.4%) | 5 mismatches/996 bases (0.5%) | 0 mismatches/996 bases (0%) | CC1 or CC80 | CC1 |
| ***sirA*** | 0 mismatches/993 bases (0%) | 6 mismatches/993 bases (0.6%) | 9 mismatches/993 bases (0.91%) | CC80 | CC1 ? |
| ***sbnA*** | 0 mismatches/981 bases (0%) | 1 mismatches/981 bases (0.1%) | 0 mismatches/981 bases (0%) | *Related in all lineages in question* | *Related in all lineages in question* |
| ***sbnB*** | 1 mismatches/1011 bases (0.1%) | 2 mismatches/1011 bases (0.2%) | 1 mismatches/1011 bases (0.1%) | CC1 or CC80 | CC1 |
| ***sbnC*** | 4 mismatches/1755 bases (0.23%) | 9 mismatches/1755 bases (0.51%) | 0 mismatches/1755 bases (0%) | CC80 | CC1 |
| ***sbnD*** | 3 mismatches/1257 bases (0.24%) | 0 mismatches/1257 bases (0%) | 3 mismatches/1257 bases (0.24%) | CC1 or CC80 | CC1 |
| ***sbnE*** | 16 mismatches/1737 bases (0.92%) | 17 mismatches/1737 bases (0.98%) | 8 mismatches/1737 bases (0.46%) | CC1 or CC80 | CC1 |
| ***sbnF*** | 0 mismatches/1740 bases (0%) | 8 mismatches/1740 bases (0.46%) | 13 mismatches/1740 bases (0.75%) | CC80 | CC1 ? |
| ***sbnG*** | 4 mismatches/777 bases (0.51%) | 4 mismatches/777 bases (0.51%) | 1 mismatches/777 bases (0.13%) | CC1 or CC80 | CC1 |
| ***sbnH*** | 5 mismatches/1203 bases (0.42%) | 3 mismatches/1203 bases (0.25%) | 2 mismatches/1203 bases (0.17%) | CC1 or CC80 | CC1 |
| ***sbnI*** | 3 mismatches/765 bases (0.39%) | 4 mismatches/765 bases (0.52%) | 2 mismatches/765 bases (0.26%) | CC80 | CC1 |
| **Q5HJP4** | *Absent from ANR570100 and 11819-97* | *Absent from ANR570100 but present in 02T-671* | 1 mismatches/408 bases (0.25%) | CC80 | CC1 |

Deviating Region 1 consists of four different fragments. The first comprises the genes between *walJ* (MS7_0024; SACOL0023) and *orfX*.

The second one is an ACME-III element including the *opp* operon. This is a potentially motile element and thus it is not necessarily connected to the genomic replacement in this strain. It will be discussed below.

A third fragment includes, among other genes, the enterotoxin homologue ORF-CM14. It extends to Q7A890/Q2YUT2 (MS7_0086/MS7_0087; SACOL0076/SACOL0077). This fragment does not contain the enterotoxin H gene *seh* or a *seh*-derived pseudogene (MS7_0080) which are characteristic for CC1 or CC80, respectively.

A forth fragment of Deviating Region 1 includes the gene encoding staphylococcal protein A. It can be assigned to RIDOM *spa* type t1849; 07-23-34-33-13. This *spa* type is related, but not identical, to both, those of CC1 (such as t127; 07-23-21-16-34-33-13) and CC80 (such as t044; 07-23-12-34-34-33-34). The RIDOM database shows 10 matches (https://spa.ridom.de/spa-t1849.shtml; as of 2020, April 3rd), including three belonging to a German project on characterisation of African *S. aureus* isolates [29, 35, 36] but without disclosing their MLST types or further details. Several other genes in this fragment match CC1 sequences (see Table 1).

### The SCC element as part of Deviating Region 1

Deviating Region 1 also comprises *orfX* together with integrated SCC elements. The reference sequence CP003194 contains a SCC*mec* IVc element and most published isolates and sequences of CC80 harbour SCC*mec* IVa or IVc elements. These are absent from ANR570100. Instead, it carries an SCC element without *mecA/C* genes.

The gene content of the SCC element is summarized in Table 2. In short, it consists of

**Table 2:** The ACME-III element in ANR570100 and 02T-671.

| Gene | Description/gene product and comments | Orientation | Start position in SCC | End position in SCC | Start position in genome | End position in genome | Comparison of ANR570100 to 02T-671 |
| --- | --- | --- | --- | --- | --- | --- | --- |
| *orfX* | 23S rRNA methyltransferase with the SCC integration site being located at the 3' end of *orfX*. | Forward | 1 | 480 | 33682 | 34162 | 0 mismatches/480 bases (0%) |
| *sRNA6* | Antisense RNA associated with *orfX*. |  | 181 | 464 | 33862 | 34146 | 0 mismatches/284 bases (0%) |
| **DR_SCC** | Direct repeat of SCC, 19 nt of the 3' end of the coding sequence of *orfX*. |  | 462 | 480 | 34143 | 34162 | 0 mismatches/19 bases (0%) |
| *dam5* | Type II restriction-modification system, endonuclease and methyltransferase. A reference sequence for this gene is from strain K12S0375, GenBank JYGF01000026.1 [127750:130506]. | Forward | 775 | 3522 | 34456 | 37204 | 0 mismatches/2748 bases (0%) |
| **helicase** | DNA helicase, associated with *dam*, putative restriction system. A reference sequence for this gene is from strain K12S0375, GenBank JYGF01000026.1 [130502:132466]. | Forward | 3512 | 5477 | 37193 | 39159 | 0 mismatches/1966 bases (0%) |
| **"YeeC"** | YeeC-like protein. 1381/1442(96%) identities and 4/1442 gaps compared to strain FORC_090, GenBank CP029198.1:39824-41262 (FORC090_0030) | Reverse | 5480 | 6920 | 39161 | 40602 | 0 mismatches/1441 bases (0%) |
| **A9UFT0** | LPXTG protein homologue. A reference sequence for this gene is from strain C427-ST42, GenBank ACSQ01000048.1 [74:295]. | Reverse | 6685 | 6906 | 40366 | 40588 | 0 mismatches/222 bases (0%) |
| **Q9KX75** | Putative protein, encoded on SCC elements. A reference sequence for this gene is from strain C427-ST42, GenBank ACSQ01000048.1 [310:813]. | Reverse | 6921 | 7423 | 40602 | 41105 | 0 mismatches/503 bases (0%) |
| **Q7A207** | Putative protein, encoded on SCC elements. A reference sequence for this gene is from strain C427-ST42, GenBank ACSQ01000048.1 [829:1143]. | Reverse | 7439 | 7750 | 41120 | 41432 | 0 mismatches/312 bases (0%) |
| **Q7A206** | Putative protein, encoded on SCC elements. A reference sequence for this gene is from strain C427-ST42, GenBank ACSQ01000048.1 [1230:1580]. |  | 7752 | 7838 | 41433 | 41520 | 0 mismatches/87 bases (0%) |
| *ccrB-1* | Cassette chromosome recombinase B, type 1. A reference sequence for this gene is from strain JCSC6690, GenBank AB705452.1 [7432:9057:r]. | Reverse | 8654 | 10282 | 42335 | 43964 | 0 mismatches/1629 bases (0%) |
| *ccrA-1* | Cassette chromosome recombinase A, type 1. 1268/1352 (94%) identities and 4/1352 gaps compared to strain JCSC6690 SCCmec type IX, GenBank AB705452.1 [10431-10962]. | Reverse | 10303 | 11652 | 43984 | 45334 | 0 mismatches/1350 bases (0%) |
| **ORF No KK12** | 513/532 (96%) identities and 1/532 gaps compared to strain JCSC6690 SCCmec type IX, GenBank AB705452.1 [10431-10962]. |  | 11655 | 12185 | 45336 | 45867 | 0 mismatches/531 bases (0%) |
| *cch* | Cassette chromosome helicase. | Reverse | 12188 | 13974 | 45868 | 47656 | 0 mismatches/1788 bases (0%) |
| **orf7795** | Putative protein from ACME element of strain C427. A reference sequence for this gene is from strain C427-ST42, GenBank ACSQ01000048.1 [7795:9354]. | Reverse | 14398 | 15957 | 48079 | 49639 | 0 mismatches/1560 bases (0%) |
| **IR_IS431** | Inverted repeat of IS431. |  | 16224 | 16239 | 49905 | 49921 | 0 mismatches/16 bases (0%) |
| **tnpIS431-06** | Transposase for IS431. | Reverse | 16284 | 16958 | 49965 | 50640 | 0 mismatches/675 bases (0%) |
| **"DLJ55_14705"** | Hypothetical protein. 97% identity compared to, GenBank CP029627.1:2809799-2816653, strain MOK042; 98% identity compared to CP029650.1:1401-8447, a plasmid from strain AR_0471, GenBank. | Forward | 17243 | 24273 | 50924 | 57955 | Cannot be assessed because of discrepancies between Nanopore and Illumina sequences. |
| **IR_IS431** | Inverted repeat of IS431. |  | 24603 | 24618 | 58284 | 58300 | 0 mismatches/16 bases (0%) |
| **F8WKF9** | Putative membrane protein. A reference sequence for this gene is from strain C427-ST42, GenBank ACSQ01000050 [548:900]. | Forward | 24821 | 25173 | 58502 | 58855 | 0 mismatches/353 bases (0%) |
| **EHQ67276** | Putative protein, branched-chain amino acid transport domain protein. A reference sequence for this gene is from strain C427-ST42, GenBank ACSQ01000050 [1158:1583]. | Forward | 25431 | 25856 | 59112 | 59538 | 0 mismatches/426 bases (0%) |
| ***opp3A/A8YYZ6*** | Putative S-adenosyl-L-methionine-dependent methyltransferase from SCC elements. A reference sequence for this gene is from strain FPR3757, GenBank CP000255.1 [77130:77945]. | Forward | 26265 | 26903 | 59946 | 60585 | 0 mismatches/639 bases (0%) |
| ***opp3B*** | Nickel/peptide ABC superfamily ATP binding cassette transporter, membrane protein | Forward | 28456 | 29412 | 62137 | 63094 | 0 mismatches/957 bases (0%) |
| ***opp3C*** | Oligopeptide permease, channel-forming protein | Forward | 29412 | 30179 | 63093 | 63861 | 0 mismatches/768 bases (0%) |
| ***opp3D/A8YZ00*** | Nickel/peptide ABC superfamily ATP binding cassette transporter, ABC protein, known from ACME elements. | Forward | 30146 | 30913 | 63827 | 64595 | 0 mismatches/768 bases (0%) |
| ***opp3E/A8YZ01*** | Nickel/peptide ABC superfamily ATP binding cassette transporter, ABC protein, known from ACME elements. | Forward | 30907 | 31540 | 64587 | 65222 | 0 mismatches/635 bases (0%) |
| **tnp_A8YYY6** | Transposase. |  | 31627 | 32293 | 65308 | 65975 | 0 mismatches/667 bases (0%) |
| **DR_SCC** | Direct repeat of SCC. |  | 33683 | 33701 | 67364 | 67383 | 0 mismatches/19 bases (0%) |

- a type II restriction-modification system,
- *ccrA/B1* recombinase genes and adjacent genes showing some similarity or relationship to SCC*mec* IX sequences (strain JCSC6690, GenBank AB705452.1),
- a large gene with repetitive sequences that is very similar to the gene encoding a hypothetical protein DLJ55_14705 in the chromosomal DNA of strain MOK042 (GenBank CP029627.1) as well as on a plasmid of a ST508/CC45 strain, AR_0471 (chromosome CP029652.1, plasmid CP029650.1)
- and an oligopeptide permease operon, *i.e., opp* genes or ACME-III as well as some genes for “putative proteins” as known from the ST42 strain C427, GenBank ACSQ.

A search of the short read archive of GenBank revealed two near-identical sequences of deviant CC80 strains, one of which (SAMEA48342418) lacked ACME-III while the other one (SAMEA3671725) harboured it, indicating a variable presence of this element in CC80 [ORF-CM14+] strains. When performing a BLAST search with the NCBI GenBank, no significant hits over the entire length of the SCC element were obtained indicating that this element has not yet been observed, although most of its genes have already been found in other SCC elements.

### Identification and characterisation of the ST567 isolate 02T-671 as a potential donor of Deviating Region 1

The observation of the enterotoxin homologue ORF-CM14 rather than of the enterotoxin H gene *seh* normally present in canonical CC1 strains, followed by a set of CC1-like genes strongly suggests that Deviating Region 1 is of chimeric origin itself. Our database of ca. 25,000 microarray hybridization profiles was searched for potential donors of Deviating Region 1, *i.e*., for strains that are chimeras harbouring ORF-CM14 in an otherwise CC1-like core genomic backbone. Only one isolate, 02T-671 a deviant, ST567/CC1 (MLST profile 10-1-1-1-1-1-1, *spa* type, t1242; 07-23-12-34-34-16-34-33-13) strain matched these criteria. However, since no genome sequence was available yet for that strain the isolate that was typed by microarray based-assays was also sequenced using Illumina Miseq.

02T-671 was a PVL-positive CC1 MSSA that differed from canonical CC1 in several features including a presence of the ORF-CM14 enterotoxin homologue and an absence of *seh*. Other differences compared to canonical CC1 strains were the presence of deviant alleles of *aur* and *isaB* as well as an absence of *cna* and Q2G1R6/*cstB* (BA000033.2: 66419-67753). It also harboured an ACME-III element (*opp* genes and *ccrAB1*). The MLST marker *arcC* was different compared to ST1 (*arcC* 10 instead of *arcC* 1) but this difference is due to a single nucleotide polymorphism indicating mutation rather than recombination.

These observations are consistent with integration of a large alien insert around *oriC.* Excluding the ACME-III element, this insert can therefore be estimated to comprise roughly 150,000 nt, ranging from between *arcC* and *aur* across *oriC* and *orfX* to Q7A890/Q2YUT2 (see Figure 1).

Deviating Region 1 of ANR570100 and the corresponding region in the ST567 isolate 02T-671 can be considered identical. This includes the gene content and the gene sequences, the presence and sequence of an ACME-III element and the fault line separating a region of unknown origin from CC1-like sequences between Q7A890 and Q2YUT2.

The ACME-III elements of ANR570100 and 02T-671 were largely identical to each other in both, gene content and gene sequences (see Table 2).

Therefore, we assume the lineage or strain represented by isolate 02T-671 to be the donor of Deviating Region 1 in the lineage of ANR570100. However, the lineage of 02T-671 is itself of chimeric nature comprising a large insert of DNA from a yet unidentified donor into a CC1 genome.

### Deviating Region 2

Deviating Region 2 (Table 3, Figure 1) extended from *htsB* (MS7_2199, SACOL2166) to Q8NVB9 (MS7_2323, SACOL2297) or to *ecfA2* (MS7_2242; SACOL2211) having a size of 33,939 to 38,645 nt (1.2 to 1.4% of the genome, which is clearly smaller than the corresponding fragment of the CC80 reference sequence which encompasses as much as 115,604 nt). The reason is that it spans the integration site that in canonical CC80 harbours a motile genomic element comprising of *hsdS, hsdM, etD*, F3TKB7, *edinB* and F5W4×2 (MS7_2226 to MS7_2231). This element is absent from ANR570100. It is also absent from all CC1 sequences. This region also comprises a gene cluster from *rplQ* (MS7_2243; SACOL2212) to *rpsJ* (MS7_2271; SACOL2240) encoding several ribosomal proteins. These genes are conserved at a very high degree in all sequences of *S. aureus* so that there was not enough variation for meaningful SNP analysis. However, when BLASTing (https://blast.ncbi.nlm.nih.gov/Blast.cgi) contig 16 (that entirely is a part of Deviating Region 2; the others are on 21 and 19), the five highest scoring matches over the entire length of the query sequence (68,165 nt) are CC1 genomes (with, e.g., 23 nt mismatches and 2 nt gaps for BX571857.1). In general, this region in ANR570100 is more related to CC1 than to canonical CC80 sequences. It also appears to be closer to 02T-671 than to MW2 but given the overall similarity of all sequences concerned, this is hard to assess. The adjacent regions, up- and downstream of Deviating Region 2, are very similar in ANR570100, 02T-671 as well as reference CC1 and CC80 sequences.

**Table 3:**
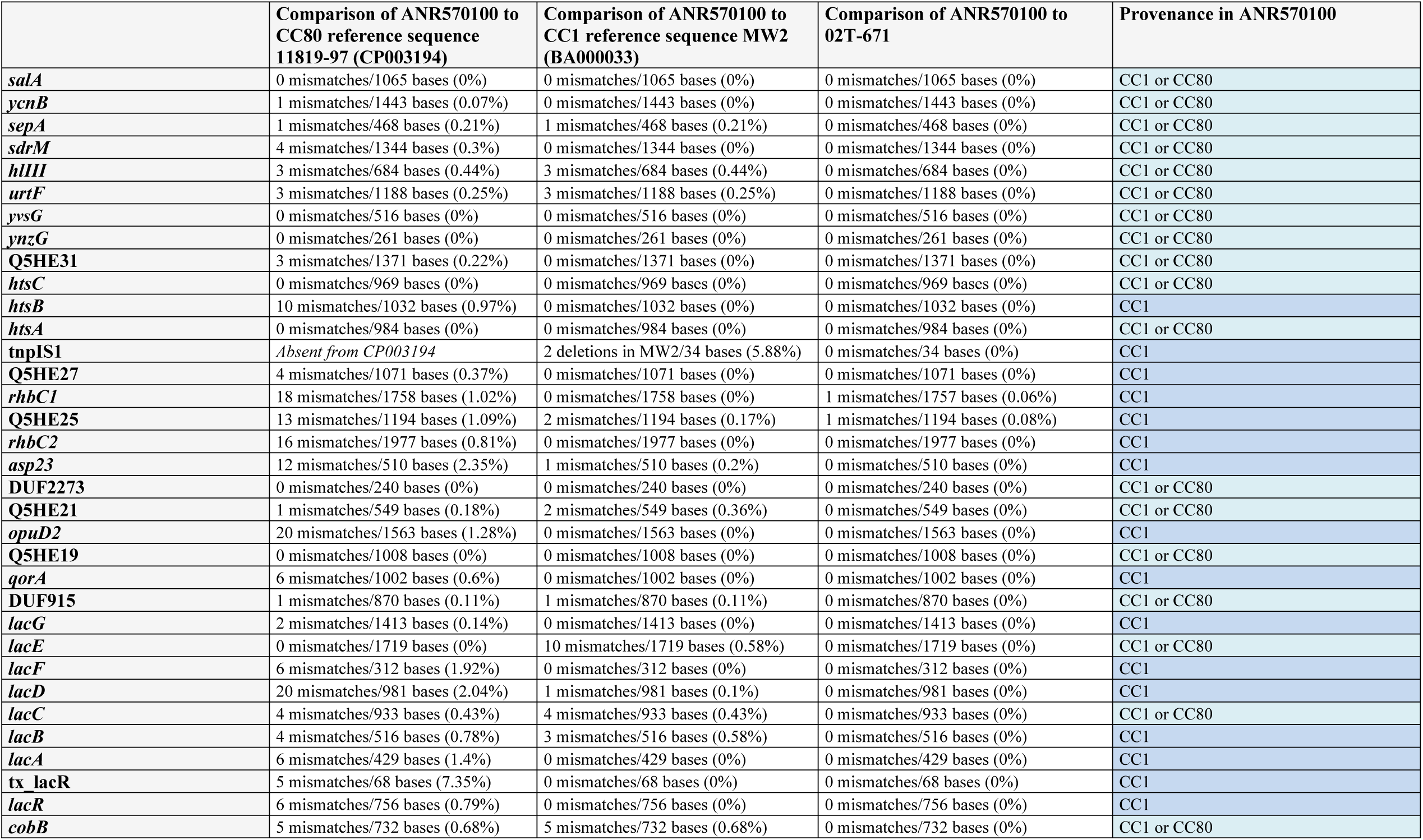
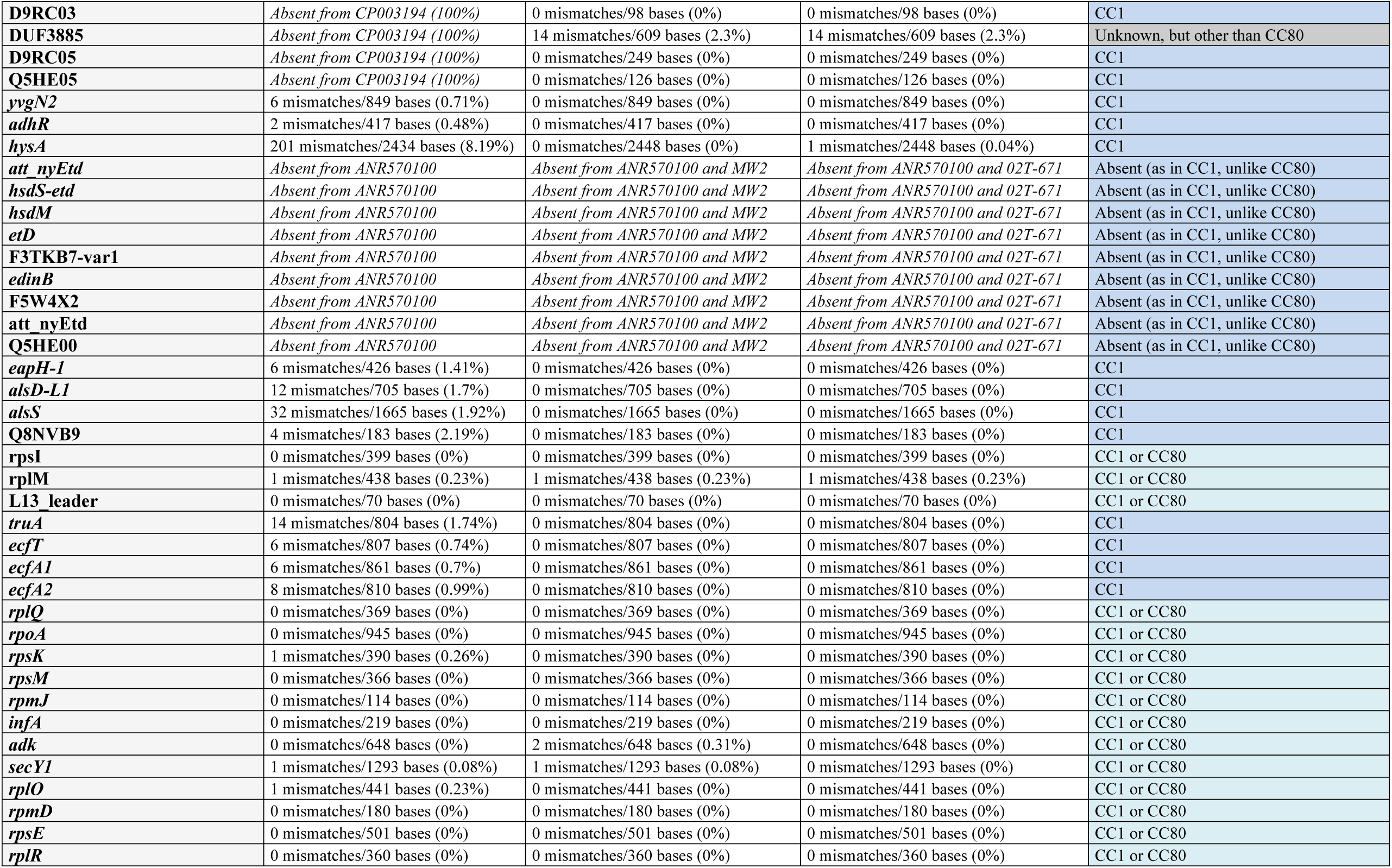
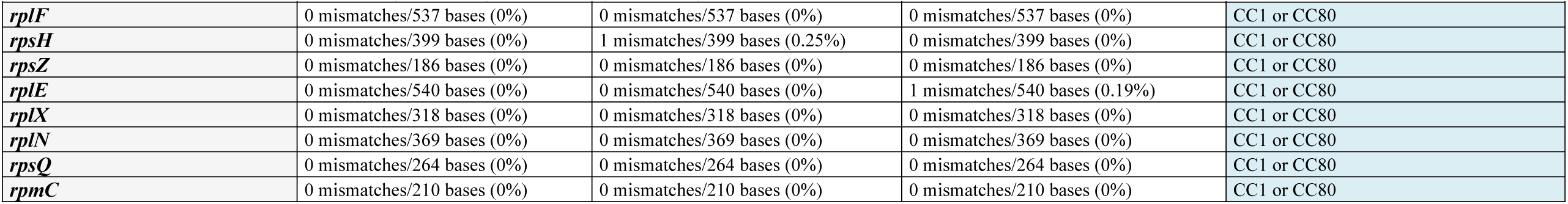
Genes in Deviating Region 2 in comparison to canonical CC80, to 02T-671 and to canonical CC1.

### The *hla* gene and its neighbouring genes

When comparing the sequence as well as the hybridization profile of ANR570100 to the CC80 reference sequence, the absence of the *hla* gene and its neighbouring genes (A5IS45, Q6GHS5, A5IS47, A6U0Y3, Q2FZB4, *i.e.*, MS7_1116 to MS7_1120 or SACOL1171 to SACOL1175) can be detected. This is presumably the result of homologous recombination between extended repeat sequences flanking the *hla* gene cluster at both of its ends. The presence of *hla* appears to be variable in the deviant CC80 lineage; SAMEA48342418 also lacks *hla* while it is present in SAMEA3671725.

### Prophages

When excising the phage sequence (Contig-0007:RC, positions 133,678 to end and Contig-0012 positions 1 to 42,938) and performing a NCBI Blast search, the four best matches, with identities of 99.97%, are all PVL phages from CC80 strains, phiSa2wa_st80 (MG029515.1), NCTC13435 (LN831036.1), GR2 (CP010402.1) and 11819-97 (CP003194.1).

The PVL prophage in ANR570100 is integrated into the same site of the chromosome as the one in 11819-97 (CP003194.1). The prophage sequences from both strains are co-linear and they comprise the same set of genes.

The *hlb*-converting phage in ANR570100 differed from CP003194.1, although the virulence-associated genes it carried (*scn, sak*) were the same. A NCBI Blast of Contig-0009RC positions 81,223 to 123,488 (as of August 2019) yielded as best matches (with more than 99.6% identity) the *hlb*-converting phages from BB155 (LN854556.1) and 55-99-44 (CP024998.1) and SA17_S6 (CP010941.1). They all belong to ST152 (https://pubmlst.org/bigsdb?db=pubmlst_saureus_seqdef&page=sequenceQuery). This might be attributed to a co-existence and co-evolution of the deviant CC80 lineage and of CC152 in the same geographic region, as the latter CC is known to be predominant at least in parts of Africa [38-44].

### Resistance genes

ANR570100 carried the *blaZ/I/R* operon and a cadmium resistance operon *cadD/cadX*, presumably on a plasmid. Genes *aphA3* and *sat* (neomycin and streptothricin resistance) as well as *far1/fusB* and *tet*(K) that frequently can be encountered in canonical CC80, being situated within SCC*mec* elements or on plasmids, were absent.

## Discussion

We identified a virulent, PVL-positive CC80 MSSA which differed in several key features from canonical CC80 strains and sequences. Its analysis was performed using three different methods, array hybridisation, Illumina and Nanopore sequencing. While array hybridisation yields less information than sequencing, it can be routinely performed automatically and economically on high numbers of clinical isolates that, in the present case, allowed the identification of the initial isolate ANR570100 as being of special interest as well as of 02T671 as putative donor. Illumina sequencing provided short reads of high quality sequences, but it has difficulties with repetitive sequences which, as the most relevant problem in the current project, led to a virtual miss of DLJ55_14705 within the ACME-III element. Nanopore sequencing proved unreliable with regard especially to poly-T and poly-A sequences, but it can handle repetitive sequences much better which in *S. aureus* also include MSCRAMM genes such as *spa*.

Differences of the target strain to reference sequences of CC80 include two large inserts of DNA from other *S. aureus* lineages, both combined accounting for about 8% of the genome of the strain. While one was located close to *oriC* which appears to be a hotspot for chromosomal replacements (see Introduction), the other one was localised at a distant position. The mechanism for these gene transfers is yet unknown. With two large replacements being present in one single isolate, we assume that such-large scale horizontal gene transfers might be more common in *S. aureus* than previously perceived, and that the resolution of MLST with seven markers is not high enough to identify all chimeric strains. However, the combination and interaction of microarray-based assays and NGS allows the reliable identification of such strains [37].

The most striking features of ANR570100, however, are large regions in its genome that clearly differ from other CC80 sequences. As described above, Deviating Region 1 in the isolate ANR570100 comprises sequences identical to the ones from the atypical CC1/ST567 strain 02T671. This includes an ACME-III element. It also includes a stretch of DNA upstream and downstream of ACME-III with the latter part including ORF-CM14. Theoretically, this might give a hint on the putative donor of Deviating Region 1.

Possible donors for ORF-CM14 to both, 570100 and 02T671 obviously must include strains form ORF-CM14 positive lineages that are ST12, ST71, ST93, ST121, ST509, ST567, CC772, CC705, ST707, ST760, ST816, ST848, ST1094, ST1643, ST2272, ST2425, ST2616 and ST2972 (based on published sequences and author’s own microarray data). Unfortunately, genome sequences of several of these STs are not available and those that are available do not match fully the sequence of Deviating Region 1. When comparing ORF-CM14 sequences alone, those of JKD6159; CP002114.2 [76914-77693] (ST93) and SS-015; FQIU01000002 [597790-598569] (ST2972) are the most closely related ones. When performing BLAST on the non-CC1-region of 02T671, the highest scoring hits are two ST2272 sequences (AP019712.1 and AP019713.1). When directly comparing sequences in question, the differences are large enough to indicate that ST2272 was not likely to be the direct donor (with an average difference of 1.8% for *dnaA, dnaN, yaaA, recF, gyrB, gyrA, nnrD, hutH, serS, azlC, sam-L1, metX, yybS, gdpP, rplI, dnaC, purA, walR, walK, walH, walI, walJ, sasH*, Q6GKL1, Q6GKL6, ORF-CM14, *dusC*, A6TXM6, A6QD71, Q6GKK6 and Q2YUT2 from Tokyo12482, GenBank AP019713.1, to 02T671).

In both strains, ANR570100 and 02T671, a fault-line can be observed between Q7A890 and Q2YUT2 separating downstream sequences of unknown origin from those upstream that are rather unambiguously related to CC1 (*i.e*., the right border between “red” and “blue” sectors in Figure 1 and the last two columns of Table 1). This means Deviating Region 1 of ANR570100 includes the fault line separating the alien insert in 02T671 from the canonical CC1 core genome of that strain. This makes it very likely that a 02T671-like strain was indeed the donor of Deviating Region 1, and that this region itself is of chimeric nature, spanning CC1 and non-CC1 sequences that together form the 02T671-like donor strain as well as the mobile SCC/ACME-III element (see Figure 1). Unfortunately, the upstream fault line separating CC1 from CC80 sequences in ANR570100 (between *sirB* and *spa* or *sbnE*) cannot exactly been determined because of the general similarity or relatedness of CC1 and CC80.

Deviating Region 1 also comprises an apparently new ACME-III element. The presence of *opp* genes and *ccrA/B-1* recombinase genes are reminiscent of the CC34 strain 21342 (GenBank AHKU) although the sequence of *ccrB*-1 appears to be more related to the one from SCC*mec* IX. It also includes, as revealed mainly by Nanopore sequencing, a gene with repetitive sequences that is very similar to the gene encoding a hypothetical protein DLJ55_14705 in strain MOK042. This strain belongs to ST71, a lineage that also can be described as chimera, comprising of a large insert of unknown origin in a CC97 genome. In strain MOK042, the gene encoding DLJ55_14705 is localised on that insert but it is not a part of a SCC element.

In addition, there is a second Deviant Region elsewhere in the genome of the lineage of ANR570100. Its gene content as well as its gene sequences are highly similar to 02T-671 and to reference CC1 sequences but they clearly differ from canonical CC80. These differences include, but are not limited to, the absence of *edinB* and *etD*. Unfortunately, the region in question includes genes whose origin cannot be determined because of a high degree of conservation of the genes affected. For the same reason, the exact boundaries of the Deviating Region cannot be identified. Interestingly, the adjacent regions to Deviant Region 2 are very similar in all sequences analysed, *i.e*., ANR570100, 02T-671 as well as the reference CC1 and CC80 sequences (with differences being less than 0.5%). This could suggest that the region corresponding to Deviant Region 2 was “deviant” not in ANR570100 but, compared to the other three sequences, in the CC80 reference sequence. This might indicate that Deviant Region 2 in ANR570100 was not an alien insert of CC1 origin but that its sequence represents shared, ancestral CC1/CC80 stock and that the corresponding region in canonical CC80 (including *edinB* and *etD*) itself was an insert from another, yet unidentified, lineage.

In conclusion, the core genome of ANR570100 bears evidence of at least two, possibly three large-scale recombination events. First, ORF-CM14, among other genes, was introduced into a CC1 strain and, second, the resulting ORF-CM14/CC1 composite fragment was introduced into CC80. In addition, another recombination event introduced either Deviating Region 2 from CC1 into the ancestor of ANR570100 or the corresponding region, possibly together with *edinB* and *etD*, from an unknown donor into canonical CC80.

Thus, such complex and large-scale recombination events cannot be that rare and exceptional, despite a mainly clonal nature of *S. aureus* [45]. Although the exact mechanism is not clear, chimerism seems to be an additional pathway in the evolution of *S. aureus*, possibly being responsible for a transmission of virulence factors (such as ORF-CM14 in the case described herein) or of resistance genes including entire SCC*mec* elements [3]. From a more theoretical point of view, large-scale genomic substitutions, chimerism or hybridisation facilitate evolutionary leaps that cannot be achieved by accumulation of single point mutations or that would require immeasurably much more time to be achieved by mutations. If one considers the ability to evolve and adapt as an evolutionary advantage, an organism that can shuffle, swap or exchange major parts of its genome by whatever unknown mechanism should be in a better position than a strictly clonal organism.

## Acknowledgements

A part of this work was presented during a poster presentation at the 18^th^ International Symposium on Staphylococci and Staphylococcal Infections (ISSSI, 23 – 26 August, 2018 Copenhagen, Denmark).

We thank for the excellent technical assistance of Byrgit Hofmann, Friedrich-Loeffler-Institut, Jena, Germany, as well as Peter Slickers, Jena, for help and advice regarding sequence analyses.

## Authors’ contributions

A. Ruppelt-Lorz and B. Söderquist found and identified strains of interest. P. Slickers, R. Ehricht and S. Monecke designed the study. A. Ruppelt-Lorz, E. Müller, D. Gawlik and A. Reißig carried out experiments; S. Monecke analysed the sequence data; H. Hotzel, S. Braun and B. Söderquist performed/supervised sequencing; A. Ziegler-Cordts created a software tool used for sequence analysis; D. Gawlik, R. Ehricht and S. Monecke wrote the manuscript. All authors read and approved the final manuscript.

## Funding

There was no external funding for this study.

## Competing interests

DG is employee of PTC - Phage Technology Center GmbH, Bönen, Germany; AZC is employee of T-Systems Multimedia Solutions GmbH, Dresden, Germany. In both cases, work on this project was performed before the respective employments started. Thus, employers of the authors did not have any role in the study design, data collection and analysis, decision to publish, or preparation of the manuscript. The other authors declare that no competing interests exist.

The authors adhere to PLOS ONE policies on sharing data and materials. The specific roles of authors are articulated in the ‘author contributions’ section.

## Availability of data and materials

The genome sequences of ANR570100 and 02T-671 are published in GenBank under the following accession numbers: *submission pending*

## Ethics approval and consent to participate

Not applicable.

## Tables, Figures and Supplemental Files

**Supplemental file 1**: Array hybridization patterns of strains discussed.

**Supplemental file 2A:** Illumina and Nanopore Consensus sequence of ANR570100.

**Supplemental file 2B:** Annotated sequence of ANR570100.

**Supplemental file 3A:** Illumina sequence of 02T671.

**Supplemental file 3B:** Annotated sequence of 02T671.

**Supplemental file 4:** fasta-file with the ACME III sequences and markers.

